# Sex-Specific Changes in Cardiac Function and Electrophysiology During Progression of Adenine-Induced Chronic Kidney Disease in Mice

**DOI:** 10.1101/2024.07.18.604201

**Authors:** Valentina Dargam, Anet Sanchez, Aashiya Kolengaden, Yency Perez, Rebekah Arias, Ana M Valentin Cabrera, Daniel Chaparro, Christopher Tarafa, Alexandra Coba, Nathan Yapaolo, Emily A. Todd, Monique M. Williams, Lina A. Shehadeh, Joshua D. Hutcheson

## Abstract

**BACKGROUND:** Chronic kidney disease (CKD) and cardiovascular disease (CVD) often co-exist and interact; however, notable sex-dependent differences are observed in how these conditions manifest and progress in parallel, despite men and women sharing similar risk factors. Identifying sex-specific diagnostic markers of cardiac structure and function throughout CKD progression could elucidate why the development and progression of these diseases differ by sex.

**METHODS AND RESULTS:** Adult, C57BL/6J male and female mice were subjected to a high-adenine (0.2%) diet throughout 12-weeks to induce CKD. Control mice were fed a normal chow diet. Every three weeks, electrocardiogram (ECG) and echocardiogram-based markers of cardiac physiology were evaluated. Adenine-induced CKD showed markers of left ventricular (LV) hypertrophy in male mice only. CKD males had markers indicative of LV systolic and diastolic dysfunction throughout regimen duration, worsening as disease progressed. Adenine males had a prolonged QTc and STc intervals when compared to Adenine females and Control males. Sex-dependent differences in the duration of the Speak-J marker, measured via ECG, was identified, with Adenine males showing increases in duration earlier than Adenine females compared to their Control counterparts.

**CONCLUSIONS:** In this study, we identified sex-dependent differences in cardiac structure, function, and electrophysiology in a mouse model of CKD-induced CVD throughout disease progression. We found that male mice are more prone to developing LV hypertrophy, systolic dysfunction, and diastolic dysfunction, with significant increases in ECG markers indicative of ventricular dysfunction observed in adenine-treated males at late stages of the disease. Additionally, we identified a new ECG parameter, Speak-J duration, that highlights sex-specific cardiac electrophysiological changes, demonstrating the model’s utility in studying sex-dependent cardiac differences.

## INTRODUCTION

Chronic kidney disease (CKD) associates with notable sex-dependent differences in cardiovascular disease (CVD). A recent study of adults with CKD found that women, when compared to men, have a lower incidence of atherosclerotic events, heart failure (HF), CVD mortality, and mortality from non-CVD causes^1^. However, women present with CKD more frequently than men^2^ and, at end-stages of CKD, experience higher rates of CVD events related to HF and stroke compared to men^3^. These sex-dependent differences in the risk of HF exist despite men and women having similar incidence of traditional risk factors, including obesity and hypertension^4^. Understanding why the development and progression of disease differ by sex, despite sharing similar risk factors, is crucial for developing targeted strategies for preventing and managing cardiovascular disease in CKD patients.

It is particularly important to identify and monitor patients with both CKD and CVD since the co-occurrence of both diseases associate with a higher rate of poor outcomes^5^. An electrocardiogram (ECG) is a noninvasive and rapid screening test that can identify abnormalities in cardiac function. ECG-specific markers can help identify the risk of HF and associated adverse events^6^. However, common ECG markers of CVD are often absent in CKD patients, which can negatively impact disease identification and monitoring^7^. A review conducted in 2019 by Skampardoni *et al.* determined that “conventional ECG markers are not reliable for risk stratification in the renal populations”^8^. To complicate matters, sex differences in ECGs among the general population are well known – with females exhibiting a higher heart rate (HR), shorter PR interval and QRS duration, and longer corrected QT duration^9^. Reliable and predictive sex-specific ECG markers of CKD-induced CVD are needed to identify patients requiring further diagnostics tests, management, and intervention.

The adenine-induced CKD mouse model, which involves feeding mice a high-adenine diet for several weeks, is commonly used to study CKD without invasive surgical procedures^10–12^. We previously showed cardiac structural and functional changes in adenine-induced CKD mice, with male mice exhibiting an increase in left ventricular (LV) ejection fraction, stroke volume index, and mass compared to control^13^. However, female mice were not included in the study. To our knowledge, no reports exist on potential differences in cardiac morphology and function between male and female mouse models of CKD. Studies in humans have focused on sex differences in CVD outcomes within the CKD population^14,15^, and sex-dependent changes in cardiac structure and function in CKD induced CVD has not been well studied.

In this study, we assess cardiac structural and functional changes during the progression of CKD-induced CVD using electrocardiography and echocardiography to determine whether these changes differ based on sex.

## METHODOLOGY

### Mouse Model

Mice of C57BL/6J background were bred and housed on a 12:12 hour light-dark cycle at room temperature (20-26°C). Adult mice, aged 8-10 weeks old and of both sexes, were assigned to either the Control group, representing healthy controls, or the Adenine group, which developed CKD and cardiac dysfunction (***Figure 1A***). Mice in the Control group were fed a normal chow diet (5V75 -PicoLab® Verified 75 IF, TestDiet®). Mice in the Adenine group were fed a high-adenine diet (0.2%), developed by Tani *et al.* to induce CKD^10^. The high adenine diet was custom made by increasing adenine (Adenine, 99%, Thermo Scientific Chemicals) levels of the normal chow diet to 0.2%. Mice were sacrificed 3, 6, 9, and 12 weeks after induction of CKD, with age-matched controls sacrificed at the same time points (***Figure 1A***). Echocardiogram and ECG were recorded at the same time points as described below. At experiment termination, body weight was assessed, and hearts were excised and weighed. A sample size of 8-10 mice was used for each time point, group, and sex.

**Figure 1:**
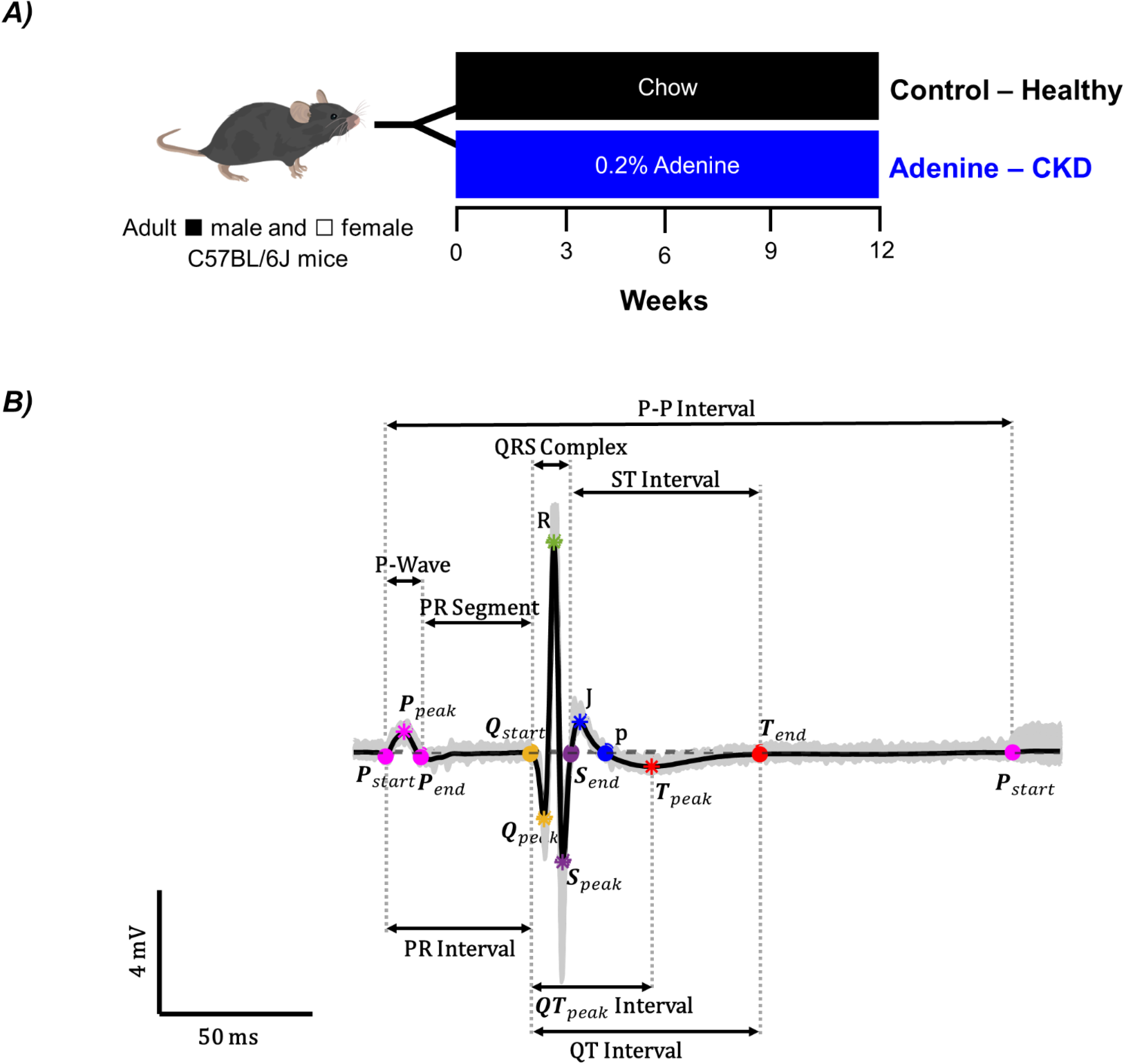
Experimental design of adenine-induced CKD and electrocardiogram (ECG) signal characteristics. ***A)*** Adult (8-10 weeks of age) C57B/6J ▪ male and □ female mice were either fed a normal chow or high adenine diet to serve as control or induce CKD, respectively. Electrocardiogram signals, echocardiograms, and tissues were collected and analyzed every three weeks starting at week 3 of the regimen. ***B)*** Mouse ECG waveform (Lead I) with its corresponding characteristics and intervals identified to quantify ECG parameters.

### Echocardiography

Cardiac hemodynamic and structural parameters were evaluated using a high-frequency ultrasound imaging system (Vevo F2, FUJIFILM VisualSonics) as previously described^13^. In short, mice we anesthetized by inhalation with an isoflurane/oxygen mixture, body temperature was maintained using a heated platform and lamp, and their physiological state (HR and respiratory rate) was monitored by recording ECG signals using subcutaneous electrodes. Cardiac parameters of structure and function were obtained using B-mode, M-mode, Pulse-Wave Doppler, and Pulse-Wave Tissue Doppler ultrasound modalities. Cardiac parameters were analyzed to evaluate overall cardiac function, systolic and diastolic function, and LV wall thickness. For measures of volume and wall size, which include ejection fraction and wall thickness, parameters were obtained in parasternal short-axis view using M-mode images. Diastolic dysfunction was assessed in apical four-chamber view, using pulse-wave doppler to quantify mitral valve blood flow parameters and pulse-wave tissue doppler to quantify mitral annular displacement. The echocardiogram data were analyzed and quantified with Vevo LAB software (FUJIFILM VisualSonics). All measurements per mouse were obtained from at least 3 cardiac cycles and the average value was used as the biological replicate for group comparison. A sample size of 6-8 mice was used for each time point, group, and sex.

### Electrocardiography

Cardiac electrical signals were recorded using Lab Rat Ephys system (Tucker-Davis Technology) and needle electrodes (29-gauge, AD Instruments). Mice were anesthetized with an inhalant anesthetic administered using a precision vaporizer. Mice were first placed in an induction chamber with isoflurane levels of 2-4% v/v and oxygen flow rate of 1-2 liters per minute. Once anesthesized, isoflurane levels of 0.5-2.5% v/v with oxygen flow rates of 1-2 liters per minute was maintained with a facemask. The ratio of oxygen to isoflurane in the inhalant anesthetic was adjusted to maintain a respiratory rate between 60-80 breaths per minute, minimizing respiratory artifacts in the ECG signals caused by deep anesthesia. Four needle electrodes were placed in the limbs subcutaneously (channel 1 to right forelimb, channel 2 to left forelimb, channel 3 to left hindlimb, and channel 4 to ground right hindlimb). ECG signals were recorded using SynapseLite (Tucker-Davis Technology) for approximately 60 seconds at a sampling frequency of 12,207 Hertz. Lead I was used for quantification of ECG parameters. A sample size of 8-10 mice was used for each time point, group, and sex. All cycles found in the recording window were used for analysis, with a minimum of 88 cycles per recording averaged to obtain each biological replicate. The location of each ECG marker was obtained from the average ECG signal.

ECG signals were processed with MATLAB (2023b, MathWorks Inc.) using a Gaussian filter over a five-element sliding window and a passband filter (5‒500 Hz) to remove high-frequency electromyographic noise. It is important to note that mouse ECG signals have different characteristics from human ECG signals^16^. For example, the QRS complex is not a good measure of total ventricular activation time in mice^16^. Therefore, markers specific to murine ECG signals were identified for this study. A custom algorithm was used to find the location of the following ECG markers: P_start_, P_peak_, P_end_, Q_start_, Q_peak_, R, S_peak_, S_end_, J, p, T_peak_, and T_end_ (***Figure 1B***). For each recording, we used an R-wave amplitude threshold to identify the location of each R-wave peak, which is indicative of a new cardiac cycle. These locations were then used to find RR interval duration. To find the remaining waves, the location of all local maxima/minima values and zero-crossings between each R-to-R segment were identified. The p marker was measured from the peak negative deflection of the Q wave to the offset of the J wave (***Figure 1B***). The offset of the J wave, or p, was defined as the beginning of negative values following the local maximum of J wave (***Figure 1B***). For mice with a J wave below zero, the S_end_ and p value were undefined, and these values were excluded from analysis. For each recording, the location of each ECG marker was blindly verified to ensure algorithm was able to accurately detect each ECG marker. In the cases where the algorithm failed to accurately detect the ECG marker, we manually selected the location.

In both mice and humans, the most common ECG parameter durations reported in literature are the following: RR interval, PR interval, P-wave duration, QRS interval, ST interval, and QT interval^17–22^. In this study, we report both traditional ECG parameters and the duration of additional, non-traditional markers in an effort to identify new ECG parameters that can detect CKD-induced cardiac abnormalities. The duration of all ECG segments (region between the end of one wave to the start of another) and intervals (region between the start of one wave to end of another) were identified (***Figure 1B***).

In humans, the duration of ECG intervals, such as the QT interval, is corrected to account for HR differences^23^. We previously found that mice on the Adenine diet have a lower HR^13^, which corresponds to a longer cardiac cycle duration. Speerschneider *et al.* showed that the QT interval duration in anesthetized C57BL/6J mice does not correlate well with HR and therefore should not be adjusted^20^. However, Boukens *et al.* argued that HR dependence could manifest in certain disease models, and ECG parameters should be corrected if a correlation exists^16^. To determine whether HR influences ECG parameters and ensure that the changes are not due to HR differences, we assessed whether a linear correlation exists between the duration of ECG characteristics to cycle duration (RR interval duration). To determine this, a linear regression model was fitted to RR interval duration and each ECG parameter, with the data separated by regimen type and combined for both groups. If the absolute value of the correlation coefficient was greater than 0.70, a strong linear relation to cycle duration was assumed and the parameter was corrected to account for HR (i.e. cycle duration). For the corrected parameter, the following formula was used to remove the linear effect of HR: *Corrected Parameter* = *Parameter* – *Slope* × [*RR Interval duration* – *Mean RR Interval Duration*]. The slope was extracted from the fitted linear model, and the mean RR Interval Duration was calculated using all sexes, regimen types, and time points.

### Tissue Collection and Morphometry

Hearts were flushed with 1xPBS by puncturing the right ventricle and flushing the circulator system with 1xPBS. Heart chambers were dissected and weighted to determine muscularization. Some hearts from mice at week 12 of the regimen were set aside for histological examination of chamber size using hematoxylin and eosin stain. Those hearts were fixed in 10% neutral buffered formalin for 24-48 hours and paraffin embedded. Transverse 5μm sections of the ventricles were stained with hematoxylin and eosin according to standard histological procedures and 2x images were obtained using a BZ-X810 microscope.

### Statistical Procedures

All parameters are presented as the mean ± standard deviation of the mean. The total number of mice per group, per time point, for each parameter is represented by n, separated by sex into females (□ F) and males (▪ M). The statistical significance of differences at each timepoint, considering sex (male or female) and regimen type (Control-Chow or CKD-Adenine), was assessed using two-way ANOVA with Bonferroni correction. A one-way ANOVA with Bonferroni correction was used to detect significance due to disease progression for each sex and its corresponding regimen type. A p-value of ≤ 0.05 was considered statistically significant.

## RESULTS

### Body Weight Affected by Regimen Type, Duration, and Sex

For both control and adenine groups, there were statistically significant differences in body weight throughout regimen duration due to regimen type and sex. For the control group, male mice were significantly heavier than female mice at all timepoints (***Figure 2B***) and both male and female mice had an increase in body weight as a function of time (***Figure 2C***). A sex-dependent difference was observed between the male and female mice in the Control group, with male mice having a higher body weight than females at each time point (***Figure 2B***). When compared to the control group, the body weight of both male and female mice in the Adenine group was significantly decreased throughout disease progression (***Figure 2C***). Male mice on the adenine group had significantly lower body weight than age-matched controls, with the adenine-fed males being 44.55% lighter than their age-matched control at week 12. Adenine-fed females were 73.57% lighter than age-matched controls at week 12. For both sexes in the adenine group, body weight decreases throughout disease progression (***Figure 2C***).

**Figure 2:**
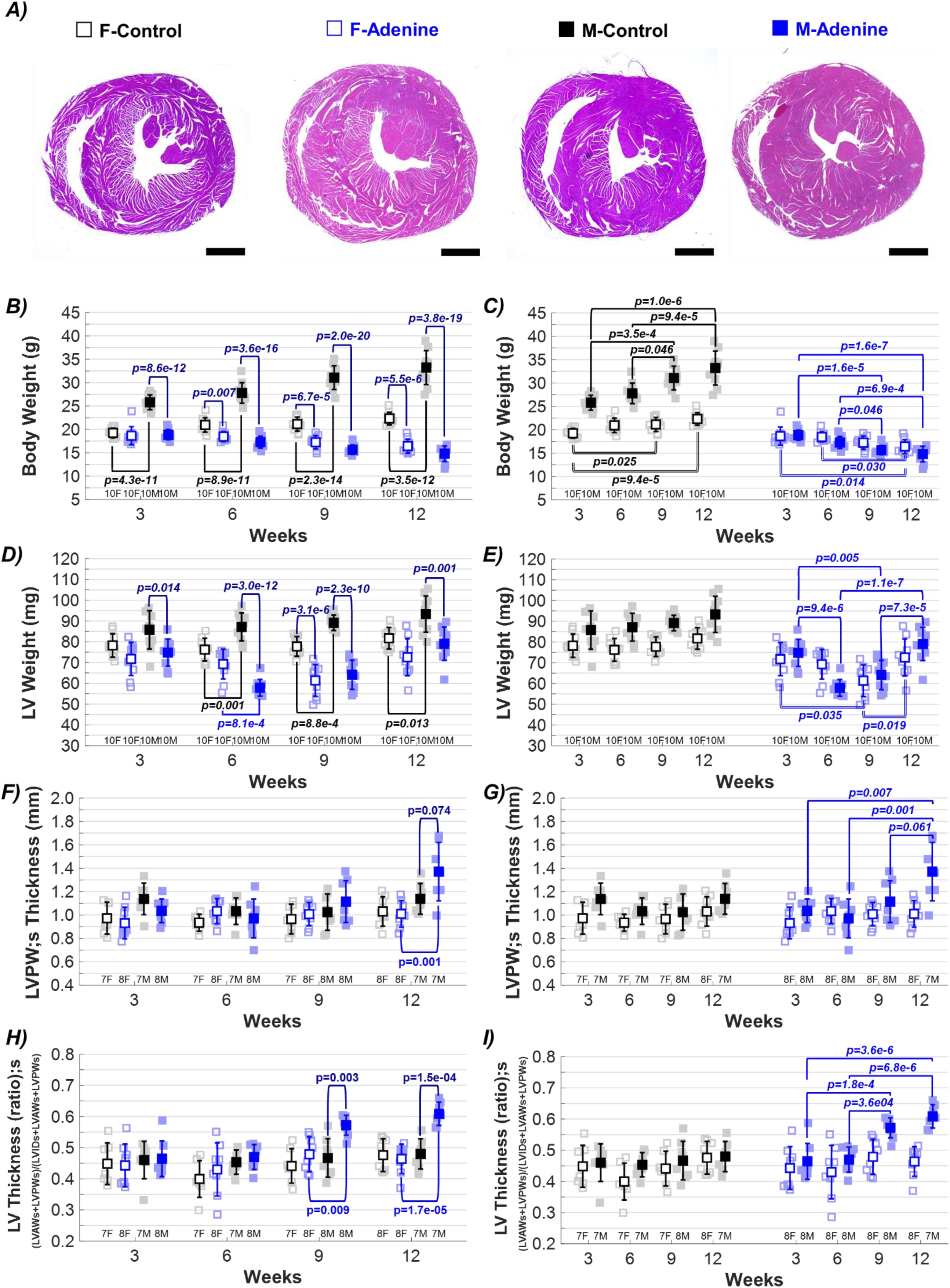
Morphological and echocardiographic evaluation of left ventricular (LV) remodeling. Changes in ▪ male and □ female mice throughout progression of either **control (Healthy)** or **adenine (CKD)** regimen. Measures of LV wall thickness and inner diameter are presented systole (s). ***A)*** Representative images of cardiac tissues using hematoxylin & eosin after 12-weeks of either dietary regimen showing size of left and right ventricles (scale bar 50 µm). Differences in ***B,C)*** left ventricular (LV) weight, ***D,E)*** left ventricular anterior wall (LVAW) and ***F,G)*** left ventricular anterior wall (LVAW) due on regiment type, duration, and sex. Results are presented as mean ± standard deviation. The statistical significance of differences at each timepoint, considering sex (▪ male and □ female) and regimen type (**Control-Chow** or **CKD-Adenine**), was assessed using two-way ANOVA (Bonferroni). A one-way ANOVA (Bonferroni) was used to detect significance due to disease progression per sex and regimen type.

### Decline in Body Weight Affects Normalization of Heart Chambers in CKD Mouse Model

Weight of heart chambers are usually normalized to either body weight or tibia length to account for variations in body size amongst animals^24^. Body weight normalization allows for a more accurate comparison of heart and chamber size relative to the overall size of the animal, providing a standardized way to assess cardiac hypertrophy. We verified the correlation between heart chambers size to body weight, heart weight, and tibia length to determine if the significant weight loss in the Adenine group and sex affected either normalization method (***Supplementary Table 1***). Heart weight and LV weight showed a linear relationship to body weight in the control group (R = 0.6609 and 0.7062, respectively). On the contrary, heart weight and LV weight correlated poorly to body weight in the Adenine group (R= 0.2051 and 0.1049, respectively). Both groups showed better linear correlations when normalizing chamber size to heart weight, with LV weight exhibiting a linear relationship in both Control and Adenine group (R = 0.8282 and 0.9077, respectively).

### CKD Induced LV Hypertrophy in Male and Female Mice

To assess cardiac hypertrophy, heart and LV weights were compared between groups and sexes. A sex-dependent difference was observed in the Control group, with male mice having larger heart weights than females at each time point (***Supplementary Table 1***). A sex-dependent difference was observed in the Adenine group at week 6 only, with male mice having smaller heart weights than females (***Supplementary Table 1***). Heart weight increased at week 9 and 12 when compared to previous time points in both male and female mice from the Adenine group (***Supplementary Table 1***). Adenine males exhibited significantly lower LV weights at weeks 3, 6, and 9 when compared to the Control group, but no difference between these groups was observed at week 12. Adenine females exhibited significantly lower LV weight when compared to Control at week 9 only (***Figure 2D***). Similar to heart weight, a sex-dependent difference in LV weight was observed between male and female Controls for weeks 6, 9, and 12 (***Figure 2D***). LV weight in Adenine mice significantly higher at week 12 compared to other time points, increasing from 64.11 mg to 79.05 mg (123.3%) in males and from 61.37 mg to 72.56 mg (110.2%) in females between weeks 9 and 12 (***Figure 2E***). A sex-dependent difference in LV weight of Adenine mice was detected at week 6 only (***Figure 2D***). No statistical differences in LV weight were identified in the control group for either male or females during the study duration (***Figure 2D-E***).

Echocardiography was also used to assess LV remodeling and hypertrophy (***Table 1***). A trending sex-dependent difference (p=0.074) in systolic left ventricular posterior wall (LVPW;s) thickness was observed at week 12 in Adenine-fed males (***Figure 2F***). Male sex and the adenine regimen were identified as factors contributing to the increase in LVPW;s size (***Figure 2F***). Adenine males demonstrated an increase in LVPW;s thickness at week 12 when compared to earlier time points, indicative of LV remodeling (***Figure 2G***). Similarly, a sex-dependent difference in the Adenine group was identified at week 12, with males showing a thicker systolic left ventricular posterior wall (LVAW;s) when compared to females but not to Control males (***Table 1***).

**Table 1.** Echocardiographic measures of cardiac function. Cardiac structural, functional, and hemodynamic changes measured via echocardiography in male and female mice fed either a **Control (Healthy)** or **Adenine** (**CKD**) diet for up to 12 weeks.

| Parameters | Week 3 |  |  |  |  |  | Week 6 |  |  |  |  |  | Week 9 |  |  |  |  |  | Week 12 |  |  |  |  |  |
| --- | --- | --- | --- | --- | --- | --- | --- | --- | --- | --- | --- | --- | --- | --- | --- | --- | --- | --- | --- | --- | --- | --- | --- | --- |
|  | Male |  |  | Female |  |  | Male |  |  | Female |  |  | Male |  |  | Female |  |  | Male |  |  | Female |  |  |
|  | Control<br>n=7M | Adenine<br>n=8M | p-value | Control<br>n=7F | Adenine<br>n=8F | p-value | Control<br>n=6-7M | Adenine<br>n=6-8M | p-value | Control<br>n=6-7F | Adenine<br>n=8F | p-value | Control<br>n=8M | Adenine<br>n=8M | p-value | Control<br>n=6-7F | Adenine<br>n=8F | p-value | Control<br>n=6-7M | Adenine<br>n=7M | p-value | Control<br>n=7-8F | Adenine<br>n=8F | p-value |
| **LV Remodeling** |  |  |  |  |  |  |  |  |  |  |  |  |  |  |  |  |  |  |  |  |  |  |  |  |
| LVAW;s (mm) | 1.28±0.11 | 1.33±0.14 | 1.0 | 1.08±0.17 | 1.15±0.16 | 1.0 | 1.25±0.11 | 1.23±0.16 | 1.0 | 1.11±0.11 | 1.17±0.15 | 1.0 | 1.26±0.16 | 1.46±0.19 | 0.113 | 0.97±0.41 | 1.21±0.14§ | 1.0 | 1.34±0.18 | 1.40±0.09 | 1.0 | 1.29±0.10 | 1.18±0.14§ | 0.475 |
| LVAW;d (mm) | 0.82±0.10 | 0.92±0.12 | 0.712 | 0.71±0.11 | 0.76±0.09 | 1.0 | 0.76±0.11 | 0.85±0.15 | 0.959 | 0.76±0.12 | 0.83±0.11 | 1.0 | 0.78±0.09 | 0.88±0.15 | 0.415 | 0.64±0.27 | 0.80±0.10 | 1.0 | 0.90±0.11 | 0.82±0.07 | 1.0 | 0.84±0.10 | 0.77±0.17 | 1.0 |
| LVPW;s (mm) | 1.14±0.14 | 1.04±0.10 | 0.812 | 0.97±0.14 | 0.93±0.14 | 1.0 | 1.03±0.11 | 0.97±0.17 | 1.0 | 0.93±0.07 | 1.03±0.11 | 0.756 | 1.02±0.15 | 1.11±0.18 | 1.0 | 0.84±0.36 | 1.01±0.10 | 1.0 | 1.14±0.13 | 1.37±0.25 | 0.074 | 1.03±0.13 | 1.01±0.11§ | 1.0 |
| LVPW;d (mm) | 0.72±0.04 | 0.70±0.06 | 1.0 | 0.68±0.10 | 0.67±0.06 | 1.0 | 0.73±0.06 | 0.64±0.10 | 0.497 | 0.69±0.05 | 0.72±0.13 | 1.0 | 0.70±0.06 | 0.76±0.14 | 1.0 | 0.60±0.25 | 0.71±0.13 | 1.0 | 0.77±0.06 | 0.80±0.06 | 1.0 | 0.73±0.07 | 0.68±0.12§ | 1.0 |
| LVID;s (mm) | 2.52±0.17 | 2.66±0.38 | 1.0 | 2.57±0.35 | 2.62±0.36 | 1.0 | 2.68±0.22 | 2.40±0.46 | 0.981 | 2.58±0.40 | 2.60±0.35 | 1.0 | 2.66±0.46 | 1.96±0.24 | 0.003 | 2.24±0.96 | 2.48±0.35§ | 1.0 | 2.65±0.34 | 1.72±0.18 | 2.2e-5 | 2.55±0.34 | 2.58±0.30§ | 1.0 |
| LVID;d (mm) | 3.83±0.13 | 3.85±0.35 | 1.0 | 3.59±0.15 | 3.69±0.29 | 1.0 | 3.96±0.16 | 3.45±0.38 | 0.009 | 3.61±0.23 | 3.68±0.26 | 1.0 | 3.94±0.30 | 3.29±0.21 | 1.1e-4 | 3.17±1.30 | 3.62±0.25 | 1.0 | 3.91±0.26 | 3.21±0.24 | 8.2e-5 | 3.70±0.23 | 3.73±0.24§ | 1.0 |
| LV Thickness;s | 0.46±0.06 | 0.47±0.06 | 1.0 | 0.45±0.07 | 0.44±0.07 | 1.0 | 0.45±0.04 | 0.47±0.04 | 1.0 | 0.40±0.06 | 0.43±0.09 | 1.0 | 0.47±0.06 | 0.57±0.03 | 0.003 | 0.44±0.06 | 0.48±0.06§ | 1.0 | 0.48±0.05 | 0.61±0.04 | 1.5e-4 | 0.48±0.05 | 0.46±0.05§ | 1.0 |
| LV Thickness;d | 0.29±0.01 | 0.30±0.03 | 1.0 | 0.28±0.02 | 0.28±0.03 | 1.0 | 0.27±0.02 | 0.30±0.04 | 0.349 | 0.29±0.03 | 0.30±0.03 | 1.0 | 0.27±0.03 | 0.33±0.03 | 0.002 | 0.28±0.02 | 0.29±0.04 | 1.0 | 0.30±0.02 | 0.34±0.02 | 0.191 | 0.30±0.03 | 0.28±0.04§ | 1.0 |
| **LV Systolic Function** |  |  |  |  |  |  |  |  |  |  |  |  |  |  |  |  |  |  |  |  |  |  |  |  |
| EF (%) | 64.0±4.0 | 59.3±6.3 | 1.0 | 55.4±10.7 | 56.6±7.8 | 1.0 | 61.0±5.9 | 59.2±8.2 | 1.0 | 55.9±11.7 | 56.9±8.3 | 1.0 | 61.5±9.9 | 72.0±6.6 | 0.112 | 49.8±21.6 | 60.3±8.3 | 1.0 | 61.0±7.2 | 79.1±3.9 | 0.001 | 59.3±9.9 | 58.9±8.3§ | 1.0 |
| FS (%) | 34.3±2.9 | 31.1±4.2 | 1.0 | 28.5±7.0 | 29.3±5.3 | 1.0 | 32.4±4.0 | 30.8±5.1 | 1.0 | 28.9±7.5 | 29.4±5.4 | 1.0 | 33.0±7.0 | 40.4±5.5 | 0.111 | 25.8±11.6 | 31.8±5.5§ | 1.0 | 32.4±5.2 | 46.6±4.1 | 3.7e-4 | 31.2±6.8 | 30.9±5.7§ | 1.0 |
| VCF<sub>c</sub> (circ/s) | 7.6±0.6 | 7.5±1.1 | 1.0 | 5.8±1.8# | 6.2±1.0 | 1.0 | 7.4±1.0 | 7.2±3.0 | 1.0 | 6.2±1.8 | 6.6±1.4 | 1.0 | 7.6±1.6 | 9.8±1.5 | 0.019 | 6.3±1.0 | 6.8±1.4§ | 1.0 | 7.5±1.2 | 12.0±2.2 | 6.9e-5 | 5.9±2.7 | 6.8±1.1§ | 1.0 |
| IVCT (ms) | 16.7±1.6 | 17.8±7.5 | 0.015 | 18.7±1.9 | 15.9±9.9 | 0.229 | 16.3±2.1 | 15.2±9.5 | 0.005 | 17.3±2.1 | 18.7±1.5 | 0.906 | 14.4±6.1 | 19.7±1.6 | 0.054 | 13.7±8.6 | 17.2±7.5 | 1.0 | 17.0±1.8 | 20.0±0.9 | 0.032 | 16.2±6.8 | 20.7±2.5 | 0.209 |
| **LV Diastolic Function** |  |  |  |  |  |  |  |  |  |  |  |  |  |  |  |  |  |  |  |  |  |  |  |  |
| MV E Velocity (mm/s) | 773.6<br>±67.2 | 669.3<br>±83.5 | 0.215 | 730.6<br>±96.3 | 736.1<br>±109.3 | 1.0 | 810.4<br>±49.5 | 605.6<br>±96.1 | 0.001 | 723.7<br>±103.5 | 683.1<br>±107.5 | 1.0 | 757.6<br>±52.5 | 627.9<br>±50.8 | 0.055 | 590.0<br>±271.0 | 680.7<br>±104.9 | 1.0 | 766.2<br>±79.6 | 667.2<br>±68.1 | 0.070 | 720.9<br>±74.1 | 720.8<br>±48.7 | 1.0 |
| MV E' Velocity (mm/s) | 23.9±5.8 | 18.5±3.6 | 0.382 | 23.7±5.5 | 25.2±6.3 | 1.0 | 22.6±3.5 | 18.1±2.8 | 0.151 | 22.5±4.4 | 22.8±3.8 | 1.0 | 19.7±1.9 | 15.8±2.3 | 0.384 | 17.2±8.1 | 24.9±5.9§ | 0.097 | 21.4±5.4 | 14.0±4.4 | 0.028 | 24.5±4.6 | 23.7±3.5§ | 1.0 |
| MV E/E' | 33.4±5.8 | 36.7±4.9 | 1.0 | 31.5±4.3 | 30.1±5.2 | 1.0 | 36.7±6.7 | 33.9±5.5 | 1.0 | 32.7±5.3 | 30.7±7.4 | 1.0 | 38.8±4.0 | 40.3±5.0 | 1.0 | 30.3±12.5 | 27.9±3.1§ | 0.012 | 37.1±6.4 | 51.2±14.6 | 0.021 | 29.9±3.6 | 30.9±4.5§ | 1.0 |
| IVRT (ms) | 24.3±2.0 | 26.1±2.4 | 1.0 | 26.2±3.7 | 27.3±4.1 | 1.0 | 18.3±8.4 | 31.2±4.0 | 1.9e-5 | 24.8±1.6 | 22.3±9.5§ | 1.0 | 22.3±2.7 | 31.2±3.5 | 5.2e-5 | 19.3±12.2 | 23.5±10.1 | 1.0 | 20.1±9.0 | 31.5±2.9 | 7.4e-5 | 21.3±9.0 | 26.7±2.5§ | 0.579 |
| **LV Overall Function** |  |  |  |  |  |  |  |  |  |  |  |  |  |  |  |  |  |  |  |  |  |  |  |  |
| Heart Rate (bpm) | 415.4<br>±40.7 | 361.3<br>±44.3 | 0.273 | 399.5<br>±42.6 | 402.3<br>±65.1 | 1.0 | 404.4<br>±31.3 | 388.6<br>±44.3 | 1.0 | 407.6<br>±52.8 | 423.0<br>±66.4 | 1.0 | 399.2<br>±26.4 | 406.0<br>±37.9 | 1.0 | 331.8<br>±142.5 | 401.4<br>±64.4 | 1.0 | 426.7<br>±24.3 | 395.1<br>±51.9 | 1.0 | 398.9<br>±52.9 | 408.9<br>±39.4 | 1.0 |
| Volume;s | 22.9±3.8 | 26.6±8.1 | 1.0 | 24.7±8.1 | 25.8±8.5 | 1.0 | 26.8±5.4 | 23.9±12.8 | 1.0 | 24.9±9.4 | 25.4±8.3 | 1.0 | 27.1±11.0 | 13.5±4.8 | 0.011 | 21.3±11.1 | 22.5±8.2 | 1.0 | 28.3±9.2 | 8.8±2.2 | 8.2e-5 | 24.1±7.8 | 24.6±6.7§ | 1.0 |
| Volume;d | 63.4±5.2 | 69.5±15.9 | 1.0 | 54.3±5.5 | 58.4±11.0 | 1.0 | 68.3±6.5 | 56.2±19.1 | 0.400 | 55.1±8.4 | 57.8±9.8 | 1.0 | 68.0±12.2 | 46.3±9.2 | 5.9e-4 | 48.6±21.1 | 55.4±9.2 | 1.0 | 71.8±17.8 | 41.7±7.5 | 1.6e-4 | 58.5±8.4 | 59.5±8.8§ | 1.0 |
| Cardiac Output (mL/min) | 16.9±2.4 | 13.6±2.5 | 0.066 | 11.9±2.3# | 13.0±2.0 | 1.0 | 16.8±1.9 | 11.1±1.4 | 3.2e-4 | 12.5±3.3# | 13.6±2.2 | 1.0 | 16.3±1.8 | 12.9±2.8 | 0.055 | 10.4±4.7# | 13.2±2.7 | 1.0 | 17.2±2.9 | 12.9±2.2 | 0.026 | 13.6±2.1 | 14.3±3.0 | 1.0 |
| Stroke Volume (µL) | 40.5±3.1 | 37.7±6.0 | 1.0 | 29.6±3.5# | 32.5±4.2 | 1.0 | 41.5±4.4 | 28.6±3.5 | 4.5e-5 | 30.3±5.4# | 32.4±4.5 | 1.0 | 40.9±4.1 | 31.6±5.2 | 9.0e-4 | 27.3±11.6# | 32.9±3.5 | 1.0 | 40.2±5.2 | 33.0±6.1 | 0.173 | 34.4±5.9 | 34.9±6.3 | 1.0 |
| ET | 54.9±3.1 | 54.5±3.4 | 1.0 | 62.3±9.8 | 57.3±5.0 | 0.661 | 53.7±4.0 | 42.2±17.7 | 0.660 | 58.8±8.8 | 53.5±5.9 | 0.692 | 53.7±4.2 | 50.3±4.8 | 1.0 | 52.2±21.3 | 60.1±8.5§ | 1.0 | 51.1±2.9 | 48.3±6.3 | 1.0 | 57.0±4.1 | 54.8±3.4§ | 1.0 |
| MPI | 0.6±0.0 | 0.7±0.1 | 0.037 | 0.6±0.1 | 0.6±0.0 | 0.049 | 0.6±0.1 | 0.7±0.3 | 3.3e-6 | 0.6±0.1 | 0.7±0.1§ | 0.003 | 0.6±0.1 | 0.8±0.1 | 0.003 | 0.5±0.2 | 0.7±0.1 | 0.324 | 0.6±0.1 | 0.8±0.1 | 8.2e-5 | 0.6±0.0 | 0.7±0.1§ | 0.481 |

Results are presented as mean ± standard deviation. Two-way ANOVA plus Bonferroni’s multiple comparisons correction used to detect significance between groups at each time point, considering regimen type and sex. If a sex-dependent interaction exists (p<0.05) at each time point, it is depicted # to the Control group and § in the Adenine group. LV thickness is defined as the ratio of LV wall size (LVAW+LVPW) to LV chamber size (LVAW+LVPW+LVID). Heart rate in this table was derived from LV trace measurements acquired in parasternal short-axis view using M-mode. Abbreviations used: **LV**, left ventricle; **s**, systole; **d**, diastole; **LVAW**, left ventricular anterior wall; **LVPW**, left ventricular anterior wall; **LVID**, left ventricular inner diameter; **EF**, ejection fraction; **FS**, fractional shortening; **VCF<sub>c</sub>**, velocity of circumferential fiber shortening, normalized to heart rate; **IVCT**, isovolumetric contraction time; **MV E**, mitral valve early flow velocity; **MV E'**, mitral annulus velocity; **IVRT**, isovolumetric relaxation time; **bpm**, beats per minute; **ET**, ejection time; **MPI**, myocardial performance index.

To account for the smaller heart size in the Adenine group, the LV wall thickness (LVAW+LVPW) was normalized to total LV size (LVAW+LVPW+LVID), defined as the sum of LV wall thicknesses (LVAW+LVPW) and LV inner diameter (LVID). Adenine males exhibited a sex-dependent difference in LV thickness ratio (systole) at week 9 and 12 (***Figure 2H***). At week 12, LV thickness in Adenine males was 120.18% greater than in Control males and 135.64% greater than in Adenine females. Additionally, LV thickness ratio in Adenine males significantly increased at week 9 and 12 when compared to previous time points of 3 and 6 weeks (***Figure 2I***).

### Male Sex Required for Manifestation of CKD-Induced LV Systolic and Diastolic Dysfunction

Echocardiography was used to assess LV systolic, diastolic, and overall cardiac function (***Table 1***). Only Adenine males exhibited signs of LV systolic dysfunction throughout disease progression, with significant differences due to regimen type and sex. Adenine males had higher LV ejection fraction (***Figure 3B***) and fractional shortening (***Figure 3D***) at week 12 compared to both age-matched Control males and Adenine females. Additionally, both parameters increase throughout disease progression and are significantly higher in Adenine males at weeks 9 and 12 compared to earlier time points (***Figure 3C, 3E***). Isovolumetric contraction time (IVCT), which reflects the period of LV contraction, was increased in Adenine males at all time points (***Figure 3H***), although not reaching statistical significance at week 9 (p=0.054). However, IVCT remained the same throughout disease progression (***Figure 3I***). Velocity of circumferential fiber shortening (VCF) is another method used to estimate LV contractile function by measuring myocardial performance^25^, and can be derived by the standard equation VCF = LV-FS/LV-ET (FS, LV Fractional Shortening; ET, LV Ejection Time). VCF is often corrected for HR to reflect the heart’s contractile function independently of rate variations. To do this, ET is corrected for HR (ETc) by dividing ET by the square root of the R-R interval (ET_c_ = ET/√R-R), with the R-R interval derived from echocardiogram-measured HR and expressed in ms. The HR-corrected velocity of circumferential fiber shortening (VCF_c_) is then calculated as VCF_c_ = LV-FS/LV-ET_c_. Similarly to EF and FS, adenine males showed a regimen and sex-dependent difference VCF_c_ at weeks 9 and 12 compared to both Control males and Adenine females (***Figure 3G***).

**Figure 3:**
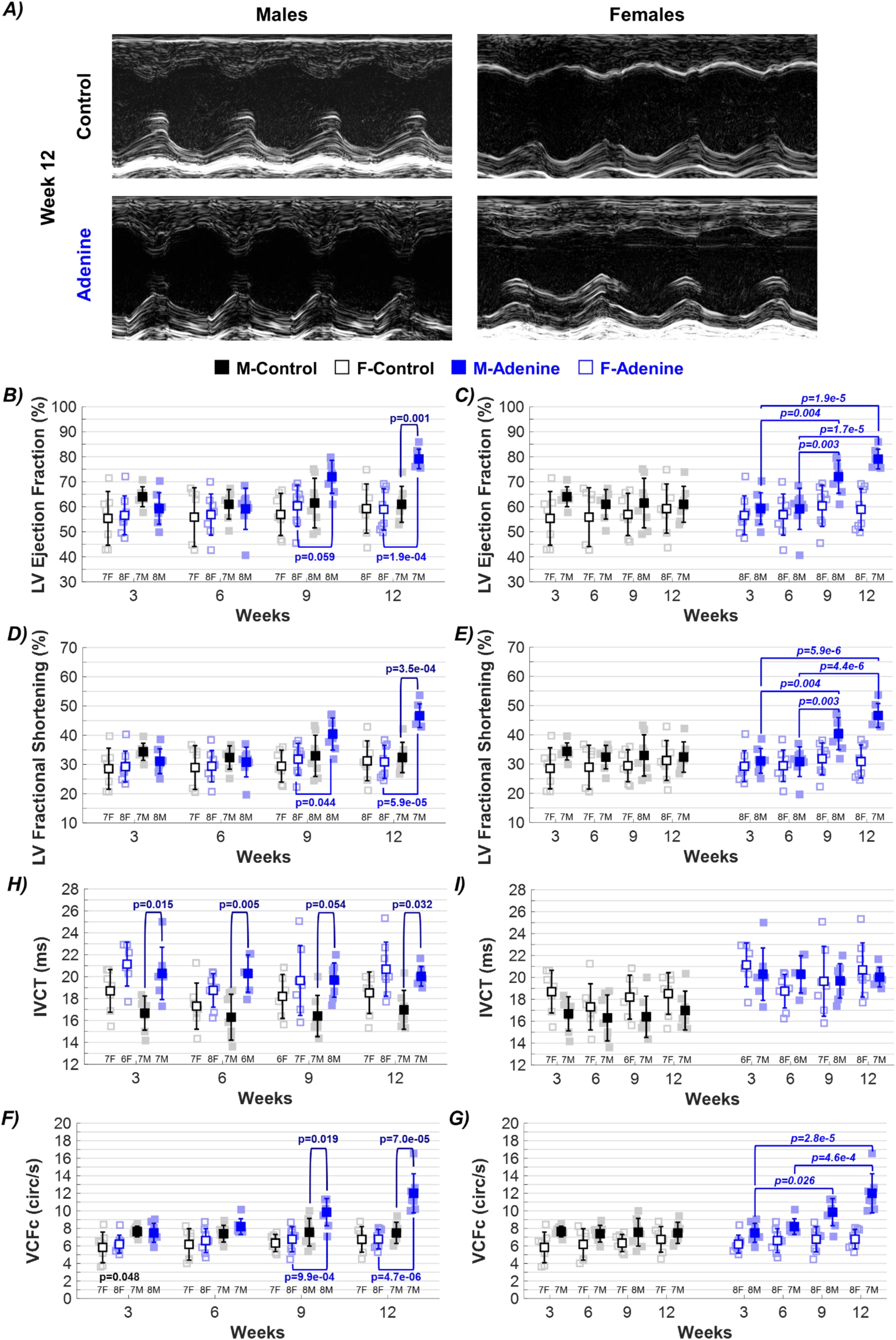
Echocardiographic evaluation of left ventricular (LV) systolic function. Changes in ▪ male and □ female mice throughout progression of either **control (Healthy)** or **adenine (CKD)** diet regimen. ***A)*** Representative M-mode images of LV chamber acquired in parasternal short-axis view. Echocardiogram-based parameters of systolic function tracked throughout disease progression included ***B,C)*** ejection fraction, ***D,E)*** fractional shortening, and ***F,G)*** isovolumetric contraction time (IVCT), Results are presented as mean ± standard deviation. The statistical significance of differences at each timepoint, considering sex (▪ male and □ female) and regimen type (**Healthy-Control** or **CKD-Adenine**), was assessed using two-way ANOVA (Bonferroni). A one-way ANOVA (Bonferroni) was used to detect significance due to disease progression per sex and regimen type.

At week 12, diastolic dysfunction associated septal mitral annulus velocity (E’) was significantly decreased (14.0±4.4 mm/s) in Adenine males compared to age-matched Control males (21.4±5.4 mm/s) and Adenine females (23.7±3.5 mm/s) (***Table 1***). Sex-dependent differences in E’ can also be observed between male and female mice in the Adenine group at week 9 (***Table 1***). Regimen- and sex-dependent differences are also observed at weeks 9 and 12 in the ratio of mitral valve early flow velocity (E) to E’, defined as the MV E/E’ ratio (***Figure 4B***). The MV E/E’ ratio gradually increased thorough disease progression in Adenine males only (***Figure 4C***). A significant increase in isovolumetric relaxation time (IVRT) was observed in Adenine males as early as week 6, which was significantly different than both Control males and Adenine females (***Figure 4D***). These changes coincide with CKD development, corresponding to the time required to observe CKD dysfunction in this mouse model^10^. An increase in MV E/E’ ratio and IVRT, as well as a decrease in MV E’ velocity, are all indicators of diastolic dysfunction in Adenine males. There was no significant difference in parameters of diastolic dysfunction in female mice due to disease manifestation nor progression at any time point (***Figure 4B-D***).

**Figure 4:**
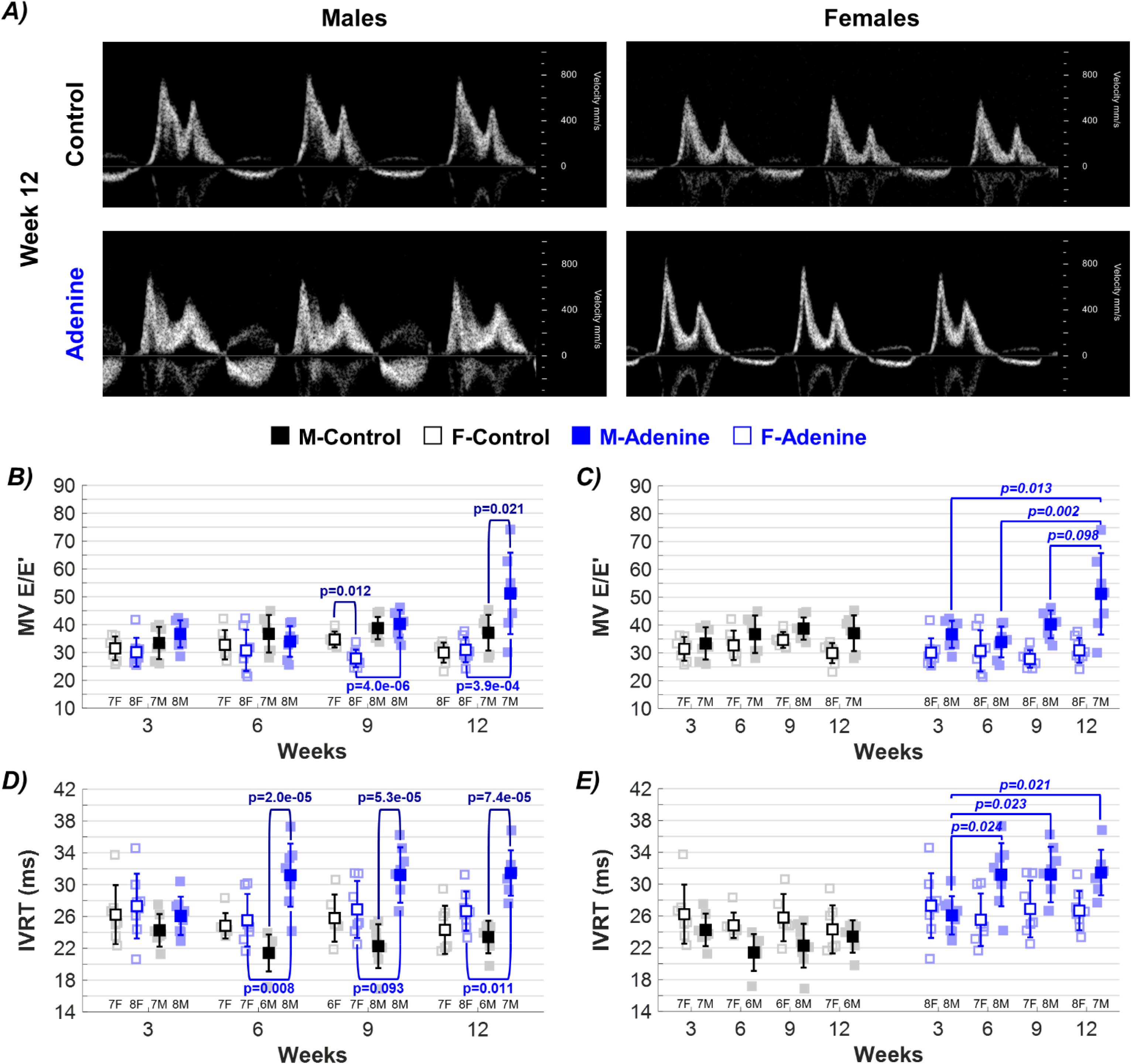
Echocardiographic evaluation of left ventricular (LV) diastolic function. Changes in in ▪ male and □ female mice throughout progression of either **control (Healthy)** or **adenine (CKD)** diet regimen. **A)** Representative Pulse-Wave Doppler mitral valve (MV) flow acquired in apical four-chamber view. Echocardiogram-based parameters of LV diastolic function tracked throughout disease progression included ***B,C)*** mitral valve early flow velocity compared to mitral annulus velocity (MV E/E’), and ***D,E)*** isovolumetric relaxation time (IVRT). Results are presented as mean ± standard deviation. The statistical significance of differences at each timepoint, considering sex (▪ male and □ female) and regimen type (**Healthy-Control** or **CKD-Adenine**), was assessed using two-way ANOVA (Bonferroni). A one-way ANOVA (Bonferroni) was used to detect significance due to disease progression per sex and regimen type.

### Cardiac Performance Exacerbated by Male Sex in CKD Mouse Model

Parameters indicative of overall LV function can be found in ***Table 1***. Adenine males had significantly lower systolic (s) and diastolic (d) LV volumes at weeks 9 and 12 when compared to Control males, with sex-dependent significant differences from Adenine females observed at week 12 (***Table 1***). Control females had a lower stroke volume than male controls at week 3, 6, and 9 (***Figure 5A***). Adenine males also exhibited a lower stroke volume when compared to Control males at weeks 6 and 9 (***Figure 5A***). Females did not show changes in stroke volume due to either regimen type (***Figure 5A***) nor over time (***Figure 5B***). Males show an increase in cardiac output throughout the regimen duration at each time point compared to females (***Figure 5C***). However, there are no changes within sex throughout regimen duration (***Figure 5D***) nor in HR (***Table 1***), indicating that changes in cardiac output differ by sex in control mice. Adenine males exhibit a decrease in cardiac output at each time point when compares to Control males but did not differ from Adenine females (***Figure 5C***). This decrease in cardiac output in Adenine males was consistent throughout the regimen duration, remaining stable despite disease progression (***Figure 5D***). Male and female mice on the Adenine group showed changes in LV myocardial performance index (MPI), an index that correlates negatively with global LV function^26^. Compared to their same-sex control counterparts, Adenine males showed a statistically significant increase in LV MPI at all time points, while Adenine females showed this increase only at weeks 3 and 6 (***Figure 5E***). There were no statistical changes in LV MPI in either sex due to disease progression (***Figure 5F***).

**Figure 5:**
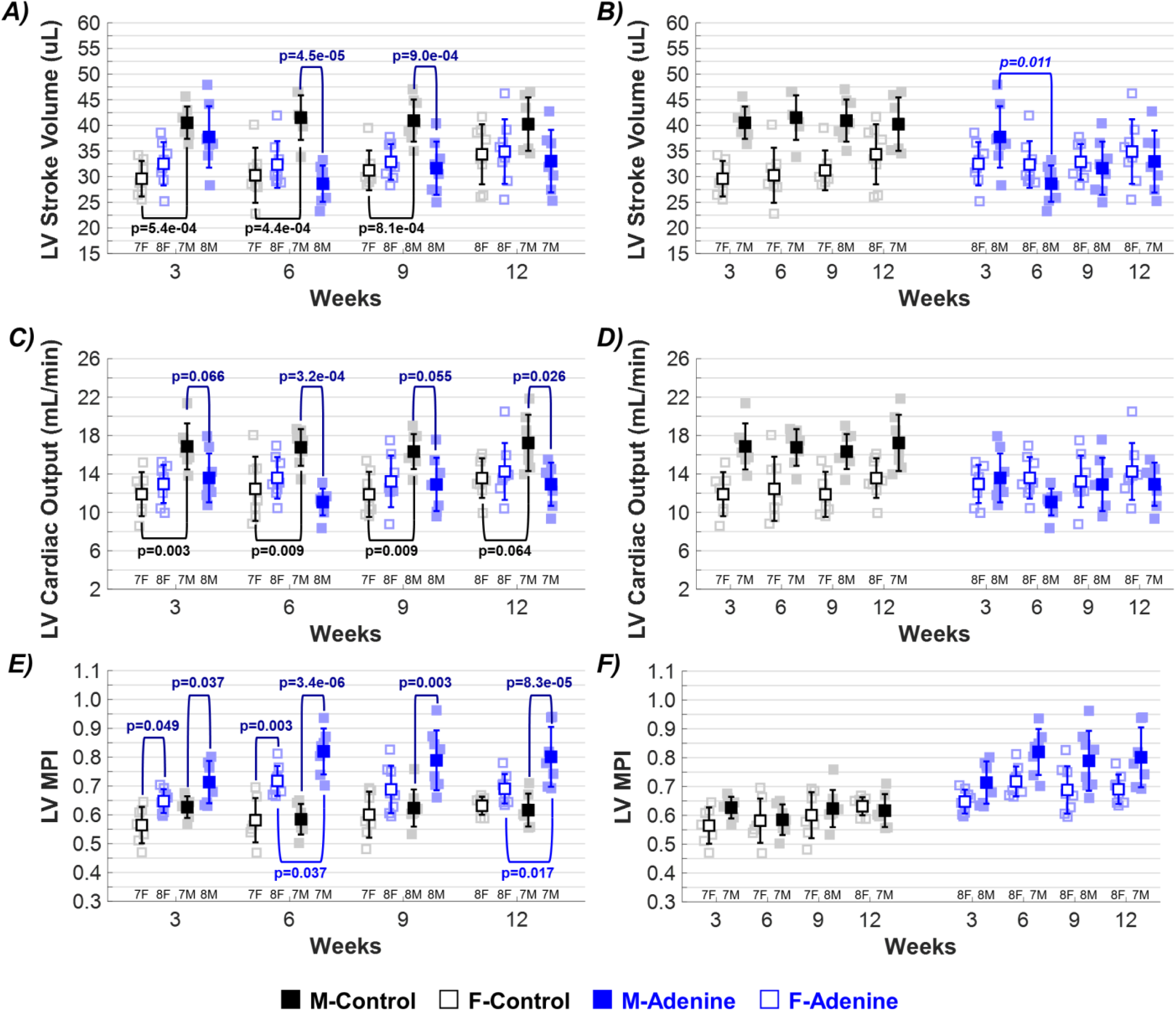
Echocardiographic evaluation of overall left ventricular (LV) function. Changes in in ▪ male and □ female mice throughout progression of either **control (Healthy)** or **adenine (CKD)** diet regimen. Echocardiogram-based parameters of overall LV function tracked throughout disease progression included ***A,B)*** stroke volume, ***C,D)*** LV cardiac output, and ***E,F)*** myocardial performance index (MPI). Results are presented as mean ± standard deviation. The statistical significance of differences at each timepoint, considering sex (▪ male and □ female) and regimen type (**Healthy-Control** or **CKD-Adenine**), was assessed using two-way ANOVA (Bonferroni). A one-way ANOVA (Bonferroni) was used to detect significance due to disease progression per sex and regimen type.

### CKD Increases Cycle Duration and Decreases HR in ECG Measurements

Representative ECG tracings are shown in ***Figure 6A*** and average duration of ECG parameters per regimen type and sex at each time point are listed in ***Table 2***. Both RR interval and PP interval durations, which measure the average duration of one cardiac cycle, are significantly increased in the Adenine groups (***Table 2***). Compared to their Control counterparts, Adenine females showed an increase in cycle duration at weeks 9 and 12, while Adenine males showed an increase at all time points (***Table 2***). A sex-dependent difference between male and female mice was only identified at week 6 of the regimen (***Table 2***). No significant difference in cycle duration due to sex or regimen duration was identified for Control mice. HR was estimated using RR and PP interval durations. Since HR is inversely correlated with cycle duration, the Adenine regimen had the same influence on HR as it did on cycle duration. Significant differences in HR estimation between groups, per sex type were present, with Adenine mice showing an increase HR at all time points compared to Controls and a sex-dependent group difference (***Table 2***). No significant difference in HR was observed due to sex or regimen duration for Control mice.

**Figure 6:**
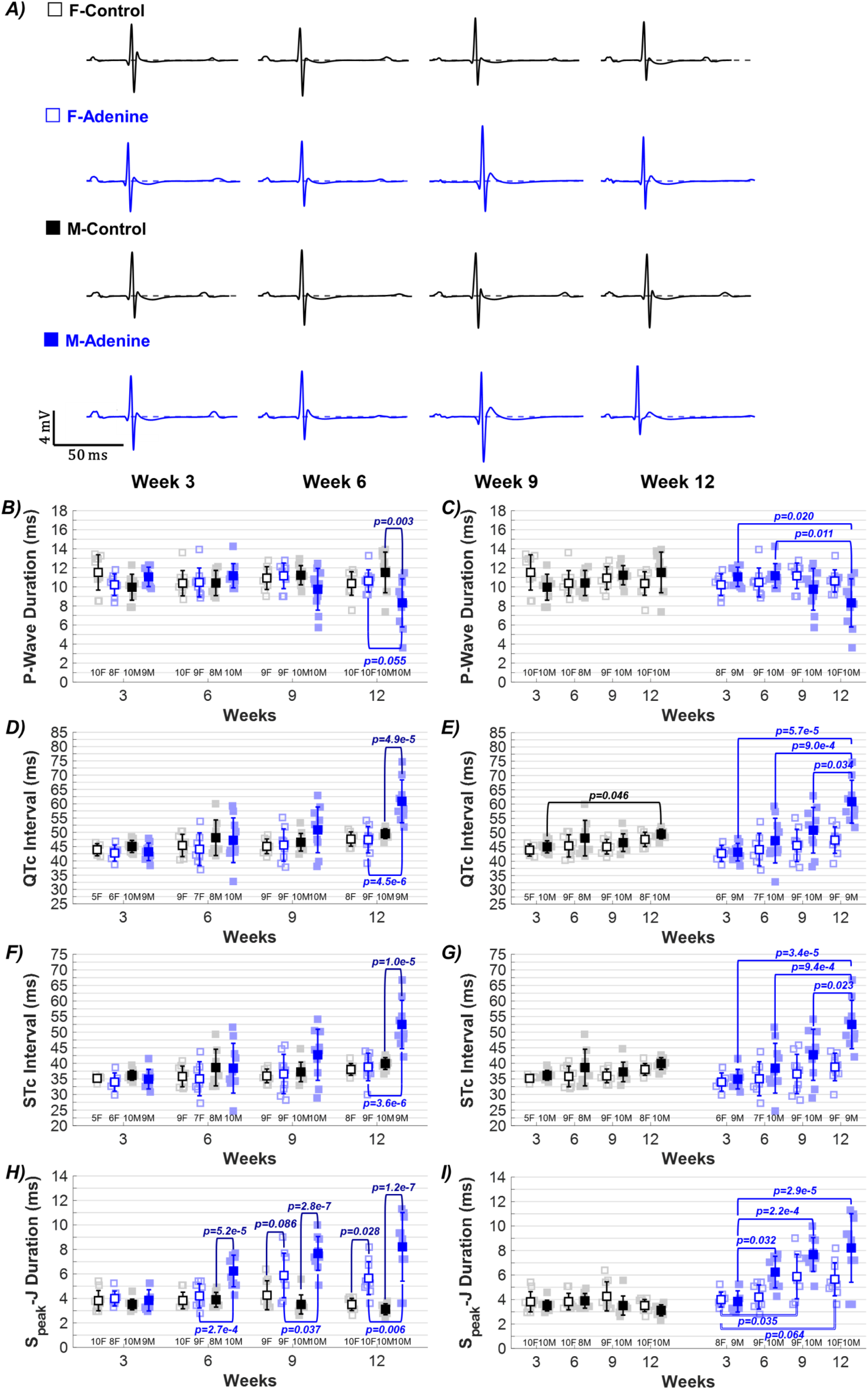
Representative electrocardiogram (ECG) signals and changes in ECG parameters due to regimen type and duration. ***A)*** Cardiac electrophysiological changes in ▪ male and □ female mice measured via ECG and quantified using Lead I throughout progression of either **control (Healthy)** or **adenine (CKD)** regimen. ECG-based parameters tracked throughout disease progression included ***B,C)*** P-wave duration, ***D,E)*** QTc interval duration, ***E,F)*** STc interval duration, and ***H,I)*** S_peak_ -J duration. Results are presented as mean ± standard deviation. The statistical significance of differences at each timepoint, considering sex (▪ male and □ female) and regimen type (**Healthy-Control** or **CKD-Adenine**), was assessed using two-way ANOVA (Bonferroni). A one-way ANOVA (Bonferroni) was used to detect significance due to disease progression per sex and regimen type.

**Table 2.**
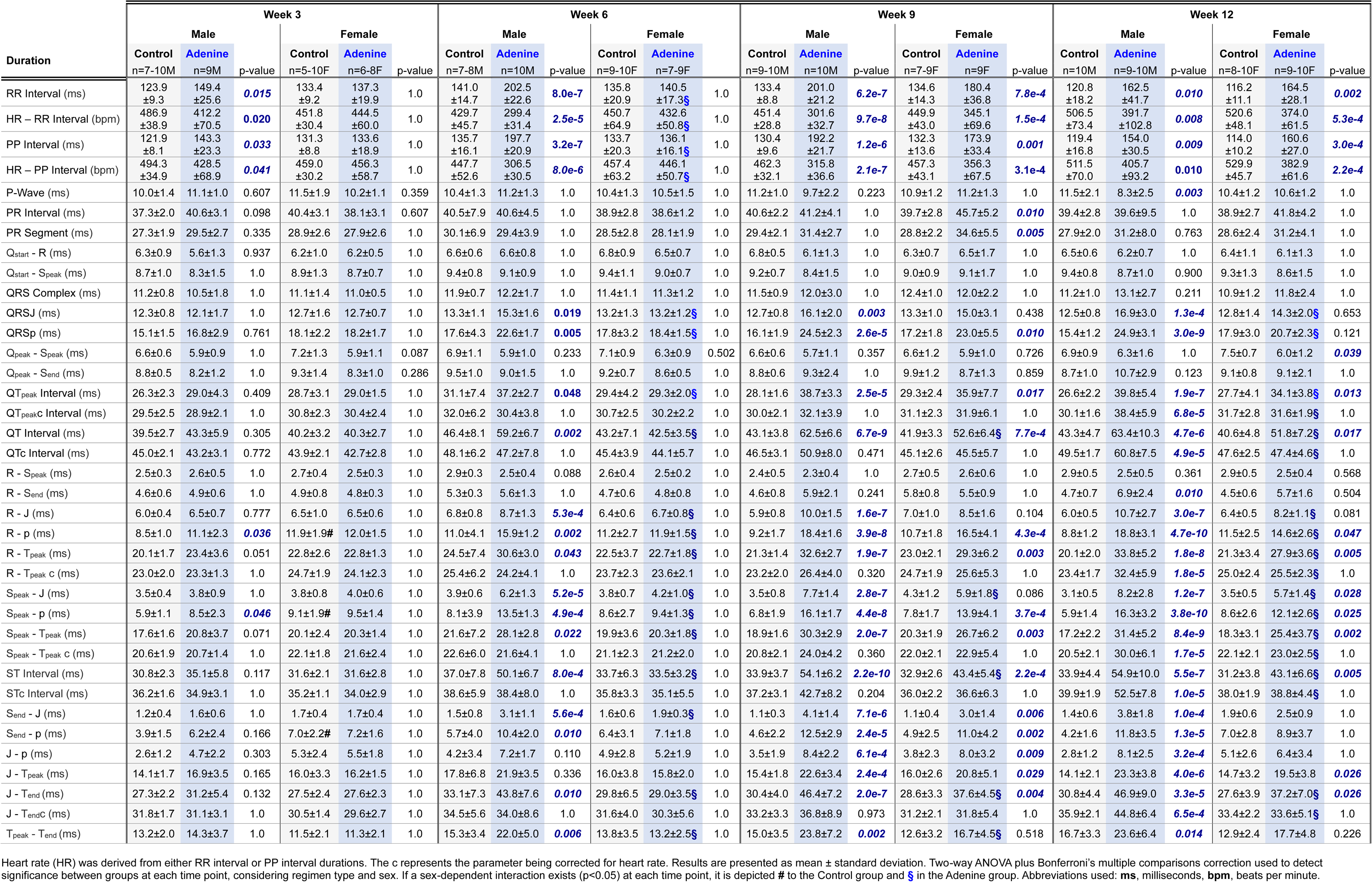
Duration of electrocardiogram (ECG) parameters. Regimen and sex-dependent differences in duration of ECG characteristics in male and female mice fed either a **Control (Healthy)** or **Adenine** (**CKD**) diet for up to 12 weeks.

### Accounting for HR Variability in the Duration of ECG Parameters

***Supplementary Table 3*** shows the correlation coefficient of all identified ECG characteristics to RR interval duration based on regimen type. The PP interval duration had a strong correlation with RR interval duration (R=0.9813), which is expected since both parameters measure the cycle duration from the start of one cycle to the next. Outside of PP interval duration, the QT interval duration had the highest correlation coefficient to RR interval duration for both the Control (R=0.7293) and Adenine groups (R=0.6552), with both groups having a combined R correlation coefficient of 0.7729 (***Supplementary Table 3***). The following parameters had an R value greater than 0.70 and were corrected (c) for cycle duration: Q-T_peak_, QT interval, R-T_peak_, S_peak_-T_peak_, ST interval, and J-T_peak_ (***Supplementary Table 3***). To ensure that HR variability was corrected for in these parameters, the linear correlation of the corrected parameters to RR interval duration was identified (***Supplementary Table 4***).

### Increases in QTc and STc Interval Durations Depend on Sex, Regimen, and HR Correction

Differences in the duration of common ECG parameters were observed due to sex and regimen type. At week 12, only the Adenine males had a significantly shorter P-wave duration (8.3±2.5 ms) compared to the Control males (11.3±2.1 ms), with no sex-dependent differences when compared to the Adenine females (***Figure 6B***). There were no statistically significant differences in PR interval duration due to regimen type at any time point, and a sex-dependent difference was only observed between Adenine males and females at week 9 (***Table 2***). The QRS complex also showed no significant differences due to sex or regimen type at any time point. Both QTc and STc interval durations show a trend of increasing throughout disease progression in Adenine males (***Figure 6D-G***), but significant differences are only observed at week 12 in comparison to Control males and Adenine females (***Figure 6D,F***). Without HR correction, an increase in QT and ST interval duration is also observed in Adenine females at week 9 and 12, with sex-dependent differences compared to Adenine males at week 12 (***Table 2***). At week 12, adenine males showed a much larger difference compared against their sex-matched Controls and Adenine females – QT duration increased by 46.37% in males and 23.82% in females, while ST duration increased by 61.95% in males and 38.14% in females.

### Sex-Specific Differences in S_peak_-J Duration Throughout CKD Disease Progression

To determine whether new ECG parameters can identify sex- and disease-dependent differences in cardiac electrophysiology, we measured the duration of various combinations of ECG markers. Of all the parameters identified, S_peak_-J duration was the only one that showed a trend throughout disease progression, with significant differences between regimen type and sex (***Figures 6H-I***). The duration of S_peak_-J differs based on sex and disease manifestation, with Adenine males exhibiting a statistically significant increase at week 6 and Adenine females at week 12 compared to Controls (***Figure H***). Adenine males were significantly different from Control males at weeks 6, 9, and 12 and Adenine females only at week 12. A strong trending difference was observed between Adenine males and females at week 9 (p=0.086) (***Figure H***). Adenine males, on average, showed an increase in S_peak_-J duration throughout disease progression, but significant differences were only identified at week 12 when compared to previous time points (***Figure 6I***). No significant differences in S_peak_-J duration were found due to sex or regimen duration for Control mice.

## DISCUSSION

Male and female sex affect physiological responses to CKD differently, influencing the progression and severity of both CKD and CKD-associated CVD^3^. Understanding how these differences influence cardiac structural, functional, and electrophysiological abnormalities can lead to more personalized approaches in managing and preventing cardiovascular complications in CKD patients. Additionally, recognizing sex-specific variations could enhance the accuracy of diagnostic tools and improve the prognostic value of existing and emerging biomarkers. This study was conducted to elucidate changes in CKD-induced cardiac dysfunction, with a particular emphasis on identifying sex-dependent variations in diagnostic CVD markers.

Cardiac morphology and function have been widely studied in the CKD-induced by 5/6 nephrectomy mouse model^17,27^. However, there are only a few studies that have assessed cardiac abnormalities in the adenine-induced CKD mouse model. Cardiac changes in this mouse model were first reported in 2018, making it a relatively new mouse model to study reno-cardiac syndrome^12^. Chen *et al.* showed that male mice on a 16-week adenine-induced CKD regimen had increased cardiac fibrotic area and increases in both LVAW and LVPW thickness, which was also observed in our mice (***Supplementary Table 2***)^28^. They also observed significant lower body and heart weights in Adenine males, similar to our findings (***Supplementary Table 2***)^12^. However, females were excluded from their study. The presence of cardiac hypertrophy in Adenine males has been well reported^12,28–30^. However, since females are mostly excluded from these studies, it is unknown whether this model also exhibits sex-dependent differences in cardiac remodeling. Our results suggest that sex-specific changes in CKD-related cardiac structural remodeling are present in this mouse model. Considering that LV hypertrophy is a common morphology that can predict the risk of adverse cardiovascular events in CKD patients, and that sex differences exist in the manifestation and progression of CKD in humans, with female sex being a protective factor in CKD-associated CVD risk, we conclude that an advantage of this model is its ability to study these CKD-induced CVD sex differences^31,32^.

A sex-dependent difference was also identified in echocardiographic-based measurements of cardiac function. Adenine males showed increases in LV ejection fraction, fractional shortening, and VCFc at week 12 of the regimen – all of which are parameters indicative of systolic dysfunction (***Figures 3B-G***). Patients with hypertrophic cardiomyopathies often exhibit an increase in ejection fraction due to maladaptive cardiac remodeling and reduced LV volume^33^. Ejection fraction, calculated as the percentage of blood ejected from the ventricle during systole relative to the total amount of blood present in the ventricle at the end of diastole, can be affected by changes in LV wall thickness and chamber size. Echocardiogram and cardiac weight dimensions showed LV hypertrophy is a sex-dependent maker of CVD in Adenine males, which had an increase in LV wall thickness and weight compared to Control males and Adenine females (***Figure 2F-I***). Although overall heart size was reduced in both Adenine males and females, as indicated by the decrease in LV weight (***Figure 2D-E***), only the LV inner diameter of Adenine males was reduced (***Table 1***). Adenine males showed a sex-dependent decrease in both systolic and diastolic LV volume at weeks 12 compared to Control males and Adenine females, leading to an increase in stroke volume (***Table 1***). Despite having a smaller LV, as measured by LV weight (***Figure 2D***), Adenine females maintained both LV volume and stroke volume (***Table 1***). We showed that changes in LV volume and wall thickness differs based on regimen type and sex in this mouse model of CKD-induced CVD, with Adenine males recapitulating clinical features of hypertrophic cardiomyopathy.

Increases in cardiac contractility have also been shown to increase ejection fraction^34^. At all regimen time points, Adenine males exhibited a prolonged IVCT when compared to Controls, but no sex-dependent differences were identified when compared to Adenine females (***Figure 3H***). Although Adenine females also had an average increase in IVCT at weeks 3, 9, and 12, IVCT was not statistically significant from Control females (***Figure 3H,I***). A prolonged IVCT strongly correlates with LV dyssynchrony, a condition that affects electromechanical activation of cardiac muscles during systole and leads to a decrease in cardiac output^35^. A prolonged IVCT is a common marker of heart failure with reduced ejection time. Heart failure with reduced ejection time and heart failure with preserved ejection fraction are both types of heart failure that can occur in CKD patients, with females more often presenting with heart failure with preserved ejection fraction^36^. In our echocardiographic data, parameters associated with diastolic dysfunction (i.e., heart failure with preserved ejection fraction) showed a male and Adenine regimen dependence. As early as week 6, Adenine males showed a significant disease- and sex-dependent increase in MV E/E’ and IVRT, both of which are common markers used to identify diastolic dysfunction in mice^37,38^. In this mouse model, the functional changes observed suggest that males experience significantly worse cardiac dysfunction compared to females. This is may be due to the fact that males exhibit more severe CKD, as previously reported by other groups^30^, thus exacerbating cardiac issues. Therefore, sex-specific differences in CKD severity are most likely a major contributing factor to the observed disparity in cardiac function in this mouse model. Since a similar sex disparity in CKD progression is observed in humans, we believe this animal model is suitable for studying sex differences in CKD-induced cardiac dysfunction.

The QTc and STc intervals were notably prolonged in adenine males at week 12, correlating with increased cardiac dysfunction. The QTc interval is a measure of cardiac depolarization and repolarization that highly correlates to ventricular arrythmias and sudden cardiac death. In CKD patients, an elongated QTc associates with higher risks of CVD events, as well as all cause and CVD mortality^39–41^. A study by King *et al.* also found a prolonged QT interval in mice fed a high (25%) and low (15%) dose dietary adenine regimen with an alternating control diet for 6 weeks^30^. More importantly, they found that changes in QT duration in CKD mice associate with severity of CKD and are sex-dependent, with Adenine males showing worse renal dysfunction and cardiac conduction than Adenine females^30^. This study, however, did not assess cardiac functional changes nor correct the QT interval for heart rate throughout disease progression. In our study, correction for HR was necessary to accurately assess sex-dependent differences in the QT interval. It has been well documented that a sex difference exists in QTc interval durations between men and women, with women showing a longer QTc than men^42–45^. Further complicating matters, changes in endogenous sex hormones throughout the menstrual cycle have been shown to influence QT interval in both humans and animals^46–48^. Age and sex specific cutoffs for prolonged QTc have been proposed to improve its prognostic value^49^. These sex-specific differences in the QT interval emphasize the importance of considering HR adjustments in ECG analysis, particularly in mouse models that significantly affect HR.

A new ECG parameter, S_peak_-J duration, showed significant sex and disease-dependent differences, especially in Adenine males. A prolonged and statistically different S_peak_-J duration was observed starting at week 6 of the regimen in Adenine males, but only at week 12 in Adenine females (***Figure 19H***). In mice, the myocardium activates early (R-wave) or late (J-wave) with a large repolarization phase, which then plateaus until the p marker and then continuously decreases until the start of the P-wave^16^. Therefore, a prolonged S_peak_-J duration is indicative of abnormal early repolarization periods. In humans, alterations in J-wave morphology are also indicative of early repolarization issues and have been linked to a higher risk of ventricular arrhythmias and sudden cardiac death^50^. Unlike in mice, a J-wave is slightly visible in humans and depends on lead placement, with a prominent J-wave appearing mostly in pathophysiological conditions^19,51,52^. Due to this, the quantification of ECG parameter durations involving this wave has not been well studied in humans. The prevalence of J-waves indicative of early repolarization has been shown to increase in CKD patients, but unlike the general population, did not seem to correlate to all-cause mortality^53^. Further studies are needed to correlate electrophysiology of the J-wave between species and assess the potential of using J-wave dependent ECG parameters to detect cardiac electrophysiological changes in CKD patients.

Although not tracked in this study, one of the disadvantages of this mouse model is a high mortality rate at later stages of disease. We observed that over 50% of the Adenine males died before reaching week 12 of the adenine regimen, while females survived over 95% of the time. Mortality rates of adenine-induced CKD mouse models in literature are contradictory. The study by Chen at al., which excluded females, found that 100% of Adenine-fed (0.20%) males survived after a 16-week regimen^28^. Kim *et al.* found that 100% of the male and female mice survived when alternating a high adenine (0.15-0.20%) and casein diet within a 10-week period^54^. Padalkar *et al.* adjusted the amount of adenine in the diet based on sex to try to reduce mortality in male mice, with males receiving an adenine dose of 0.1% and females 0.2%^55^. However, they still showed that males on an 0.1% adenine diet had a much lower survival rate than females in a 0.2% adenine diet, with ∼20-25% of males surviving compared to ∼90% of females^55^. The reported differences in mortality rates are most likely influenced by experimental design, with type of adenine used to supplement normal diet, adenine dosage, alternating between an adenine-enriched and normal diet, and regimen duration all contributing to these discrepancies.

Another important contributing factor to consider that could contribute to sex-dependent differences in mortality is the progression of both CKD and CKD-induced CVD. Our results show that male sex plays an important role in the manifestation and progression of CKD-induced CVD, with Adenine males exhibiting significant changes in structural, functional, and electrophysiological markers of disease. Adenine males had a shorter P-wave duration, which has been associated with increased risks of atrial fibrillation (***Figure 6B,C***)^56^. As previously discussed, the adenine-induced CKD mouse model shows a prolonged QTc duration with sex-dependent differences in progression of CKD, with female sex being protective against CKD^30^. The same study also showed that CKD mice had an increased risk of sudden cardiac death due to progressive bradyarrhythmias but did not specify whether this risk varied by sex^30^. We hypothesize that this sex-dependent difference in CKD-induced CVD progression also contributes to mortality rate, with Adenine males progressing in both diseases faster than females. This phenomenon is also observed in humans, with males progressing faster and having a higher incidence of end-stage kidney disease than females despite females having a higher incidence of CKD^57,58^. We conclude that this mouse model can be useful for studying sex disparities in CKD-induced CVD with the caveat regarding the need for careful considerations of study design as noted above. In addition to an increased mortality rate, Adenine males significantly lose body weight, with Adenine males being less than 50% of the size of sex-matched Controls (***Figures 2B-C***). This drastic reduction in body weight can falsely exaggerate changes in other cardiac structures if used for their normalization. To prevent this but still account for changes in body size, we tried normalizing cardiac structure to heart weight. Normalization of LV weight to heart weight only showed significant differences between Adenine and Control males at week 3 (***Supplementary Table 2***). Control males had a lower LV/Heart weight ratio at week 3 compared to other weeks (***Supplementary Table 2***). Reduction in body size, as well as progression of disease, greatly affect procedures requiring anesthesia. This also interferes with the acquisition of echocardiograms, as the ribs become more pronounced and the heart twists severely to the left as body weight is reduced, making it difficult to visualize and evaluate cardiac function. Further studies are needed to determine the best type of adenine regimen to elucidate sex-dependent differences in CKD-induced CVD progression while reducing the complications associated with this model.

## CONCLUSIONS

In this study, we identified sex-dependent differences in cardiac structure, function, and electrophysiology in an Adenine-fed mouse model of CKD-induced CVD throughout disease progression. We determined that male sex is crucial for the development of LV hypertrophy, LV systolic dysfunction, and LV diastolic dysfunction. We showed a regimen-, duration-, and sex-dependence in the duration of common ECG parameters, with Adenine males exhibiting an increase in QTc duration compared to Control males and Adenine females at late stages of the disease. cardiac functional changes correlate to ECG characteristics. More importantly, we identified a new ECG parameter (Speak-J duration) that can identify sex-specific cardiac electrophysiological changes Adenine mice. The findings in this study show the feasibility of using this mouse model of CKD-induced CVD to study sex-dependent differences in cardiac structure, function, and electrophysiology.

## ACKNOWLEDGEMENTS

None.

## REFERENCES

1. Toth-Manikowski SM, Yang W, Appel L, Chen J, Deo R, Frydrych A, Krousel-Wood M, Rahman M, Rosas SE, Sha D, et al. Sex Differences in Cardiovascular Outcomes in CKD: Findings From the CRIC Study.

2. Muhammad A, Ayesha S, Mohsin H, Haseeb F, Aastha P, Rosheen J, Siffat SS, Lateef IK, Deepak S, Rajasekar R, et al. Sex-specific differences in risk factors, comorbidities, diagnostic challenges, optimal management, and prognostic outcomes of heart failure with preserved ejection fraction: A comprehensive literature review. Heart Failure Reviews. 2023. doi: 10.1007/s10741-023-10369-4

3. Shah S, Christianson AL, Meganathan K, Leonard AC, Crews DC, Rubinstein J, Mitsnefes MM, Schauer DP, Thakar CV. Sex Differences in Cardiovascular Outcomes in Patients With Kidney Failure. Journal of the American Heart Association. 2024. doi: 10.1161/jaha.123.029691

4. Yolande A, Yolande A, Bas BvR, Bas BvR, Monique EtH, Monique EtH, Eric B, Eric B, Sanne P, Sanne AEP. Sex differences in cardiovascular risk factors and disease prevention. Atherosclerosis. 2015. doi: 10.1016/j.atherosclerosis.2015.01.027

5. David HS, Thorp M, Gurwitz J, McManus D, Goldberg R, Allen L, Grace H, Sung S, Magid D, Go A. Chronic Kidney Disease and Outcomes in Heart Failure With Preserved Versus Reduced Ejection Fraction. 2013.

6. Wesley TON, Matylda M, Alain GB, David AB, Mouaz HA-M, Joao ACL, Dalane WK, Elsayed ZS. Electrocardiographic predictors of heart failure with reduced versus preserved ejection fraction: The multi-ethnic study of atherosclerosis. Journal of the American Heart Association. 2017. doi: 10.1161/jaha.117.006023

7. Arunkumar S, Subbiah A, Arun Kumar S, Yogesh C, Yogesh KC, Sandeep M, Sandeep M. Cardiovascular disease in patients with chronic kidney disease: a neglected subgroup. Heart Asia. 2016. doi: 10.1136/heartasia-2016-010809

8. Sofia S, Sofia S, Sofia S, Dimitrios P, Dimitrios P, Dimitrios P, Marek M, Marek M, Marek M, Darren G, et al. The potential of electrocardiography for cardiac risk prediction in chronic and end-stage kidney disease. Nephrology Dialysis Transplantation. 2019. doi: 10.1093/ndt/gfy255

9. Ernst S, Ernst S, Henry B, Henry B, Thomas CP, Thomas P, Thomas CP, Pauline E, Pauline E, Fernanda R, et al. Sex Differences in the Electrocardiogram. Circulation. 1960. doi: 10.1161/01.cir.22.4.598

10. Takashi T, Hideo O, Akira S, Shuichi T. Development of a novel chronic kidney disease mouse model to evaluate the progression of hyperphosphatemia and associated mineral bone disease. Scientific Reports. 2017. doi: 10.1038/s41598-017-02351-6

11. Miki T, Mizuho T, Reiko A, Reiko A, Masaru H, Masatoshi H, Hiroshi O, Hiroshi O. Progressive renal dysfunction and macrophage infiltration in interstitial fibrosis in an adenine-induced tubulointerstitial nephritis mouse model. Histochemistry and Cell Biology. 2009. doi: 10.1007/s00418-009-0557-5

12. Julius K, Julius K, Jianmin C, Jianmin C, Samira A, Samira A, Samira A, Paul C, Paul C, Kieran M, et al. A novel model of reno-cardiac syndrome in the C57BL/ 6 mouse strain. BMC Nephrology. 2018. doi: 10.1186/s12882-018-1155-3

13. Valentina D, Hooi Hooi N, Sana N, Daniel C, Camila Iansen I, Suhas Rathna S, Armando B, Zachary CD, Lina AS, Joshua DH. S2 Heart Sound Detects Aortic Valve Calcification Independent of Hemodynamic Changes in Mice. Frontiers in Cardiovascular Medicine. 2022. doi: 10.3389/fcvm.2022.809301

14. Sultana S, Sultana S, Janaki A, Janaki A, Jacqueline KP, Jacqueline KP, Cara MH, Cara MH. Relationship between sex and cardiovascular mortality in chronic kidney disease: A systematic review and meta-analysis. PLOS ONE. 2021. doi: 10.1371/journal.pone.0254554

15. Dorothea N, Dorothea N, Josef C, Morgan EG, Morgan EG, Yingying S, Yingying S, Corri B, Corri B, Massimo C, et al. Associations of estimated glomerular filtration rate and albuminuria with mortality and renal failure by sex: a meta-analysis. BMJ. 2013. doi: 10.1136/bmj.f324

16. Bastiaan JB, Mathilde RR, Mathilde RR, Stacey R, Ruben C, Ruben C. Misinterpretation of the mouse ECG: ‘musing the waves of Mus musculus’. The Journal of Physiology. 2014. doi: 10.1113/jphysiol.2014.279380

17. Judith R, Radloff J, Nejla L, Latic N, Ulrike P, Pfeiffenberger U, Christiane S, Christiane S, Simone T, Tangermann S, et al. A phosphate and calcium-enriched diet promotes progression of 5/6-nephrectomy-induced chronic kidney disease in C57BL/6 mice. Scientific Reports. 2021. doi: 10.1038/s41598-021-94264-8

18. Polina SS, Polina S-S, Lars S, Lars S, Abraham MR, Abraham MR, Kathleen LG, Kathleen LG. Electrocardiographic Characterization of Cardiac Hypertrophy in Mice that Overexpress the ErbB2 Receptor Tyrosine Kinase. Comparative Medicine. 2015.

19. Charlotte C, Petra S. What to consider for ECG in mice—with special emphasis on telemetry. Mammalian Genome. 2023. doi: 10.1007/s00335-023-09977-0

20. Tobias S, Tobias S, Morten BT, Morten BT. Physiology and analysis of the electrocardiographic T wave in mice. Acta Physiologica. 2013. doi: 10.1111/apha.12172

21. Manuela AO, Janine MW, Shin-ichi A, Ghina Bou A, Tsz Kwan C, Jae-Hoon C, Dave C, Emily MD, Elfertak L, Alain G, et al. Comprehensive ECG reference intervals in C57BL/6N substrains provide a generalizable guide for cardiac electrophysiology studies in mice. Mammalian Genome. 2023. doi: 10.1007/s00335-023-09995-y

22. Mohamed H, Mohamed H, Mohamed H, Asmaa M, Asmaa M, Kuanquan W, Kuanquan W, Feng J, Feng J, Moussa A, et al. Detection of abnormal heart conditions based on characteristics of ECG signals. Measurement. 2018. doi: 10.1016/j.measurement.2018.05.033

23. Anand A, Anand A, Anand A, Swee-Guan T, Swee-Guan T, Abdul Razakjr O, Abdul Razakjr O, Kian-Keong P, Kian Keong P, Kian Keong P. Importance of QT interval in clinical practice. Singapore Medical Journal. 2014. doi: 10.11622/smedj.2014172

24. Quint AJH, Guido PLB, Koop AMC, Anne-Marie CK, Arnold P, Tim RE, Diederik EvdF, Herman HWS, Rudolf AdB, Rolf MFB. A novel method optimizing the normalization of cardiac parameters in small animal models: the importance of dimensional indexing. American Journal of Physiology-heart and Circulatory Physiology. 2019. doi: 10.1152/ajpheart.00182.2019

25. Wolfram HK, Wolfram HK. Relationships between mean velocity of circumferential fiber shortening (VCF) and heartrate -the diagnostic value of a normalization of VCF to heart rate. Journal of Clinical Ultrasound. 1978. doi: 10.1002/jcu.1870060105

26. Jörg S, Jörg S, Markus AE, Markus AE, Clemens T, Clemens T, Markus R, Markus R, Lars E, Lars E, et al. Echocardiographic assessment of global left ventricular function in mice. Laboratory Animals. 2009. doi: 10.1258/la.2007.06001e

27. David JK, David JK, Jihad E, Jihad E, Amjad S, Amjad S, Anna PS, Anna PS, Smaili S, Sleiman S, et al. Partial nephrectomy as a model for uremic cardiomyopathy in the mouse. American Journal of Physiology-renal Physiology. 2008. doi: 10.1152/ajprenal.00472.2007

28. Cheng C, Cheng C, Cheng C, Caidie X, Caidie X, Hanzhang W, Hanzhang W, Lin W, Lin W, Lin W, et al. Uraemic Cardiomyopathy in Different Mouse Models. Frontiers of Medicine in China. 2021. doi: 10.3389/fmed.2021.690517

29. Yinghui H, Yinghui H, Shaobo W, Shaobo W, Jie Z, Jie Z, Jie Z, Yong L, Yong L, Changhong D, et al. IRF1-mediated downregulation of PGC1α contributes to cardiorenal syndrome type 4. Nature Communications. 2020. doi: 10.1038/s41467-020-18519-0

30. Benjamin MNK, Shana M, Xianming L, Gregory EM, Florencia S, Alireza KJ, Glenn IF. Chronic Kidney Disease Induces Proarrhythmic Remodeling. Circulation: Arrhythmia and Electrophysiology. 2023. doi: 10.1161/circep.122.011466

31. Sheila KP, Elena V, Daniel G, Jay R, Jessica L, Louise MB. Left ventricular hypertrophy in experimental chronic kidney disease is associated with reduced expression of cardiac Kruppel-like factor 15. BMC Nephrology. 2018. doi: 10.1186/s12882-018-0955-9

32. Ana CR, Ana CR, Wei Y, Wei Y, Daohang S, Daohang S, Lawrence JA, Lawrence JA, Jing MC, Jing C, et al. Sex-related disparities in CKD progression. Journal of The American Society of Nephrology. 2019. doi: 10.1681/asn.2018030296

33. Marvin AK, Marvin AK, François MA, Francois MA. Ejection Fraction: Misunderstood and Overrated (Changing the Paradigm in Categorizing Heart Failure). Circulation. 2017. doi: 10.1161/circulationaha.116.025795

34. Mathew SM, Mathew SM, Mathew SM, Jonathan SB, Jonathan S-B, Lyna El-Khoury R, Lyna El-Khoury R, Madeline Y, Madeline Y, Donald LK, et al. Mechanisms Underlying Improvements in Ejection Fraction With Carvedilol in Heart Failure. Circulation-heart Failure. 2009. doi: 10.1161/circheartfailure.108.806240

35. Frederik HV, David V, Lars G, Matthias D, Maximo R-A, Philippe De V, Hugo Van H, Rozette R, Pieter MV, Pieter V, et al. Revisiting diastolic filling time as mechanistic insight for response to cardiac resynchronization therapy. Journal of the American College of Cardiology. 2013. doi: 10.1016/s0735-1097(13)60611-7

36. Ida L, Ida L, Karolina S, Karolina S, Ulf D, Ulf D, Tomas J, Tomas J, Lars HL, Lars L. Associations with and prognostic impact of chronic kidney disease in heart failure with preserved, mid-range, and reduced ejection fraction. European Journal of Heart Failure. 2017. doi: 10.1002/ejhf.821

37. María V-O, María V-O, Pablo G-P, Pablo GP, Enrique L-P, Enrique LP. Non-invasive assessment of HFpEF in mouse models: current gaps and future directions. BMC Medicine. 2022. doi: 10.1186/s12916-022-02546-3

38. Moritz S, Moritz S, Norman C, Norman C, Min Z, Min Z, Adam N, Adam AN, Anderson GA, Grace A, et al. Echocardiographic evaluation of diastolic function in mouse models of heart disease. Journal of Molecular and Cellular Cardiology. 2018. doi: 10.1016/j.yjmcc.2017.10.006

39. Rajat D, Rajat D, Haochang S, Haochang S, Elsayed ZS, Elsayed ZS, Wei Y, Wei Y, Joshua MA, Joshua MA, et al. Electrocardiographic Measures and Prediction of Cardiovascular and Noncardiovascular Death in CKD. Journal of The American Society of Nephrology. 2016. doi: 10.1681/asn.2014101045

40. Peter F, Peter F, Stephen OP, Stephen OP, Abinav G, Abinav G, William M, William WM, Rachel EP, Rachel EP. Associations of ECG interval prolongations with mortality among ESRD patients evaluated for renal transplantation. Annals of Transplantation. 2014. doi: 10.12659/aot.889927

41. Mirela D, Mirela D, Andrei B, Andrei B, Arash R, Arash R, Mahboob R, Mahboob R. Electrocardiogram Abnormalities and Cardiovascular Mortality in Elderly Patients with CKD. Clinical Journal of The American Society of Nephrology. 2012. doi: 10.2215/cjn.07440711

42. Simon WR, Simon WR. Impact of Age and Sex on QT Prolongation in Patients Receiving Psychotropics. The Canadian Journal of Psychiatry. 2015. doi: 10.1177/070674371506000502

43. Robert JG, Robert JG, James RB, James RB, Zuoyao C, Zuoyao C, Keaven MA, Keaven MA, Emanuela HL, Emanuela HL, et al. Duration of the QT interval and total and cardiovascular mortality in healthy persons (The Framingham heart study experience). American Journal of Cardiology. 1991. doi: 10.1016/0002-9149(91)90099-7

44. Guy S, Guy S, Glenna CLB, Glenna CLB. Sex differences in the mechanisms underlying long QT syndrome. American Journal of Physiology-heart and Circulatory Physiology. 2014. doi: 10.1152/ajpheart.00864.2013

45. Salama G, Bett G. Sex and Gender Differences in Cardiovascular Physiology – Back to Basics Sex differences in the mechanisms underlying long QT syndrome. 2014.

46. TomoakiSaito, AndreaCiobotaru, Jean C, LigiaToro, EnricoStefani, MansourehEghbali. Estrogen contributes to gender differences in mouse ventricular repolarization. Circulation Research. 2009. doi: 10.1161/circresaha.108.190041

47. Alan HK, Alan HK, Philip G, Philip G, Marian CL, Marian CL, William HF, William HF, Sandra AD, Sandra AD, et al. Estrogen and Progestin Use and the QT Interval in Postmenopausal Women. Annals of Noninvasive Electrocardiology. 2004. doi: 10.1111/j.1542-474x.2004.94580.x

48. Kararigas G, Nguyen BT, Jarry H. Estrogen modulates cardiac growth through an estrogen receptor α-dependent mechanism in healthy ovariectomized mice. Molecular and Cellular Endocrinology. 2014;382:909–914. doi: 10.1016/j.mce.2013.11.011

49. Ilan G, Ilan G, Arthur JM, Arthur JM, Arthur JM, Wojciech Z, Wojciech Z. QT interval: how to measure it and what is “normal”. Journal of Cardiovascular Electrophysiology. 2006. doi: 10.1111/j.1540-8167.2006.00408.x

50. Abdi A, Nida B, Azeem SS. Early repolarization syndrome: A cause of sudden cardiac death. World Journal of Cardiology. 2015. doi: 10.4330/wjc.v7.i8.466

51. Gan-Xin Y, Gan-Xin Y, Charles A, Charles A. Cellular Basis for the Electrocardiographic J Wave. Circulation. 1996. doi: 10.1161/01.cir.93.2.372

52. Sven K, Sven K, Sander V, Sander V. Cardiac electrophysiology in mice: a matter of size. Frontiers in Physiology. 2012. doi: 10.3389/fphys.2012.00345

53. Reza H, Reza H, Ronak R, Ronak R, Kaivan K, Kaivan K, Sébag F, Frederic AS, Frederic AS, Soudeh M, et al. The prevalence of electrocardiographic early repolarization in an adult cohort with chronic kidney disease and its impact upon all-cause mortality and progression to dialysis. Frontiers in Physiology. 2013. doi: 10.3389/fphys.2013.00127

54. Kyoungrae K, Kyoungrae K, Kyoungrae K, Kyoungrae K, Erik A, Erik A, Trace T, Trace T, Guanyi L, Guanyi L, et al. Skeletal myopathy in CKD: a comparison of adenine-induced nephropathy and 5/6 nephrectomy models in mice. American Journal of Physiology-renal Physiology. 2021. doi: 10.1152/ajprenal.00117.2021

55. Mugdha VP, Alexandra HT, Ylona G, Mariya K, Neil K, Danyang M, Michael CG, Kelly AB, Puneet D, Saud N, et al. Paradoxical reduction of plasma lipids and atherosclerosis in mice with adenine-induced chronic kidney disease and hypercholesterolemia. Frontiers in Cardiovascular Medicine. 2023. doi: 10.3389/fcvm.2023.1088015

56. Ian CC, Ian CC, Ian CYC, Erin A, Erin A, Balaji K, Balaji K, David GB, David GB, Chwee NQ, et al. Shorter minimum p-wave duration is associated with paroxysmal lone atrial fibrillation. Journal of Electrocardiology. 2014. doi: 10.1016/j.jelectrocard.2013.09.038

57. Kunitoshi I, Kunitoshi I, Shigeru N, Shigeru N, Takahiro S, Takahiro S, Yuji N, Yuji N, Takashi A, Takashi A. Increasing gender difference in the incidence of chronic dialysis therapy in Japan. Therapeutic Apheresis and Dialysis. 2005. doi: 10.1111/j.1744-9987.2005.00318.x

58. Manfred H, Manfred H, Brian B, Brian B, Jean É, Jean E, Alexandra KW, Alexandra K-W, Gere SP, Gere S-P, et al. Sex-Specific Differences in Hemodialysis Prevalence and Practices and the Male-to-Female Mortality Rate: The Dialysis Outcomes and Practice Patterns Study (DOPPS). PLOS Medicine. 2014. doi: 10.1371/journal.pmed.1001750

